# Mechanisms of Kinesin-1 activation by Ensconsin/MAP7 *in vivo*

**DOI:** 10.1101/325035

**Authors:** Mathieu Métivier, Brigette Y. Monroy, Emmanuel Gallaud, Renaud Caous, Aude Pascal, Laurent Richard-Parpaillon, Antoine Guichet, Kassandra M. Ori-McKenney, Régis Giet

## Abstract

Centrosome separation in *Drosophila* larval neuroblasts and asymmetric transport of embryonic determinants in oocytes are both microtubule-dependent processes that require Kinesin-1 activation by Ensconsin/microtubule-associated protein 7 (MAP7). However, the molecular mechanism used by Ensconsin to activate Kinesin-1 remains elusive. Ensconsin/ MAP7 contains an N-terminal microtubule-binding domain (MBD) and a C-terminal Kinesin-binding domain (KBD). Using rescue experiments in live flies, we show that KBD expression alone is sufficient to fully rescue Ensconsin-dependent centrosome separation defects, but not the fast oocyte streaming and the localization patterns of Staufen and Gurken proteins. Interestingly, we show here for the first time that KBD binds and stimulates Kinesin-1 binding to Mts *in vivo* and *in vitro*. We propose that the KBD/Kinesin-1 motor represents a minimal activation module that stimulates Kinesin-1 binding to Mts. Addition of the MBD, present in the full length Ensconsin allows this activation to occur directly on the Mt. Our data also suggest that in a very large cell with a complex microtubule network, but not in smaller cells, this dual activation by Ensconsin is essential for optimal Kinesin-1 targeting to the microtubule cytoskeleton.

## Introduction

Microtubules (Mts) are dynamic polymers that grow and shrink over time, a process known as dynamic instability^1^. They are regulated by microtubule-associated proteins (MAPs), which have a wide range of activities. MAPs can modify microtubule dynamic parameters, and are able to organize microtubules into complex structures. The MAP family also includes microtubule motors that slide along the Mts. These motor proteins use ATP hydrolysis to move cargo along microtubules, thus transporting vesicles, chromosomes, proteins, RNA, and even other microtubules. Mt networks and their associated MAPs therefore play key roles in various biological processes, including cell division, intracellular trafficking, and cell morphogenesis^2–4^

Ensconsin/MAP7 was first identified in microtubule preparations isolated from epithelial cells^5^. The protein has been given several names, including epithelial MAP of 115 kD (E-MAP115) and Ensconsin, due to its tenacious association with microtubules both *in vivo* and *in vitro*^5^. Ensconsin/MAP7 is associated with the interphase microtubule cytoskeleton, but its overexpression in cultured cells does not seem to affect microtubule dynamics during interphase, and instead mitotic abnormalities frequently occur during cell division^6^.

The possible function of Ensconsin remained elusive for a decade, until *ensc* mutant flies were isolated in a screen for genes affecting *Drosophila* female germ line development. *ensc* mutant females are sterile, and oocytes display microtubule-dependent mislocalization of several polarized key molecules required for the proper patterning of developing embryos. These molecules include *oskar* mRNA and its binding partner, the Staufen adaptor protein, on the posterior side^7–9^, and the Gurken protein that encodes a TGF beta like protein on the dorso-ventral axis^9, 10^. The *ensc* mutant oocytes also show slower microtubule-dependent streaming of granules^9^. Interestingly, these defects are shared by Kinesin heavy chain (KHC) mutants^11–14^. In *ensc* oocyte extracts, the recruitment of Kinesin-1 motors to Mts is impaired *in vitro*, and it has recently been shown that purified Ensconsin and MAP7 proteins can directly recruit kinesin-1 to the Mt *in vitro*, providing evidence for a model in which Ensconsin/MAP7 promotes efficient recruitment of Kinesin-1 to Mts^9^. This Ensconsin/MAP7 model of Kinesin-1 recruitment is also evident in mouse and *Drosophila* muscle fibers. In these polyploid cells, both Ensconsin and Kinesin-1 are required to promote nuclei positioning and spacing, processes required for correct muscle function. In this study, the authors also showed a direct physical association between the Kinesin-1 motor and the Ensconsin C-terminal domain (the Kinesin-binding domain, or KBD). Furthermore, after Kinesin-1 or Ensconsin knock down, a fusion between the Ensconsin N-terminal Mt-binding domain (MBD) and the Kinesin-1 motor domain is sufficient to rescue nucleus-positioning defects^15^. Together, the studies of muscle cells and fly oocytes cited here support a model in which Ensconsin favors Kinesin-1 microtubule recruitment via its MBD and KBD domains^9,15^. However, a second model, based on S2 cultured cell studies, proposes that some Ensconsin function does not depend on the MBD. For these functions, Kinesin-1 can be successfully activated solely by the KBD^16^, showing that microtubule targeting by Ensconsin is not an absolute requirement for Kinesin-1 function.

Here, we used a combination of genetic studies with rescue constructs at different stages of fly development as well as *in vitro* experiments to unambiguously show that Kinesin-1 is subjected to a dual activation by Ensconsin that synergizes its loading onto microtubules.

## Materials and Methods

### DNA constructs

Histidine-tagged expression constructs were generated as previously described using full-length Ensconsin; Ensc-MBD, the N-terminal domain which contains the microtubule-binding domain, also referred to as EHR1; and Ensc-KBD, the C-terminal domain containing the Kinesin-binding domain, also referred as EHR2^9, 17^. Ensc-MBD and Ensc-KBD were also introduced into the pUWG vector to generate fly expression constructs for the ubiquitous expression of GFP-tagged proteins. Full-length Ensconsin was introduced in pTWV vector to generate over expression of a VFP-protein under the control of the GAL protein. pUWG and pTWG were purchased from the Drosophila Genomics Resource Center (Indiana University). For *in vitro* experiments Ensconsin proteins were cloned in frame using Gibson cloning into a pET28 vector with an N-terminal strepII-Tag, superfolder GFP (sfGFP) cassette.

### Fly strains

Flies were maintained under standard conditions at 25°C. All studies were performed using *enscΔnull* and *enscΔN ensconsin* mutants^9^ In contrast with previous studies, Western blotting experiments using an antibody raised against the C-terminal region of Ensconsin revealed the *enscΔN allele* is a null allele. *enscΔnull* is also a null allele but the deletion also removes a piece of the neighboring gene^9^ All experiments were therefore performed in trans-heterozygous *enscΔnull/enscΔN* flies, hereafter referred to as *ensc* flies. Transgenic flies were obtained from BestGene (Chino Hills, CA) following P-element-mediated transformation with the pUWG vector containing the FL-Ensc, Ensc-KBD, or Ensc-MBD sequences. The Ensc-GFP and Ensc-KBD-GFP transgenic proteins rescued the viability of the *enscΔnull* and *enscΔN* mutants and the trans-heterozygous *ensc* flies, but the Ensc-MBD-GFP transgenes could not. RFP-tubulin expressing flies were given to us by Renata Basto (Institut Curie, Paris). The 69B-GAL4 strain is from Bloomington Stock Center and was used to drive over expression of Ensconsin-Venus in brain tissues (Ensc OE). All other GFP and RFP fusion proteins were ubiquitously produced under the control of the polyubiquitin promoter.

### Antibodies and western blotting

The monoclonal YL1/2 rat anti-detyrosinated tubulin antibody (1:200) and the mouse monoclonal and rabbit polyclonal anti-phosphorylated histone H3 (Ser10) antibodies (1:500) were obtained from Millipore. The mouse anti-GFP monoclonal antibody (1:1000) was obtained from Roche. The rabbit and rabbit anti-actin (1:4000) polyclonal antibodies, the anti-Staufen antibody (1:1000) came from Santa Cruz Biotechnology. The rabbit polyclonal anti-KHC antibody (1:2000) was obtained from Cytoskeleton. The mouse monoclonal anti-Gurken antibody (1:200) was from the Developmental Studies Hybridoma Bank. The making of the anti-Ensconsin antibody was previously described, and the antibody was raised against the Kinesin-binding domain^17^ The mouse anti-strepII (1:5000) antibody was from Fisher NBP243719). The mouse anti-tubulin clone DM1A (1:2500) was from Sigma.

The goat peroxidase-conjugated secondary antibodies (1:5000) were obtained from Jackson ImmunoResearch, and donkey Alexa fluor-conjugated secondary antibodies (1:1000) came from Life Technologies. For western blotting, ECL reagent was purchased from Thermo Fisher Scientific.

### Live microscopy

Brains expressing the various GFP-or RFP-tagged proteins were dissected in Schneider’s *Drosophila* medium containing 10% FCS. Isolated brains were loaded and mounted on stainless steel slides. The preparations were sealed with mineral oil (Sigma) as previously described^17^. Images were acquired using a spinning-disk system mounted on an Eclipse Ti inverted microscope (Nikon) equipped with a 60X 1.4 NA objective at 25°C. Z-series were acquired every 30 or 60 seconds with a Photometrics CCD camera (CoolSNAP HQ2) and an sCMOS ORCA-Flash4.0 (Hamamatsu) controlled by MetaMorph acquisition software (version X). Images were processed using ImageJ software and are presented as maximum-intensity projections. The mother and daughter centrosomes are easily distinguished by their different microtubule nucleation potentials measured after separation, after cytokinesis, or immediately before mitosis^18^. Embryos were collected on agar plates supplemented with grape juice at 25°C. They were dechorionated by hand using double-sided adhesive tape, then mounted in mineral oil as previously described^19^. Z-series images were acquired every 30 seconds using either a spinning disk system or a Leica SP5 confocal microscope equipped with a 40X 1.3NA objective at 25°C. For ooplasmic streaming analyzes, young females were mated with males and fattened with dried yeast for 3 (Controls, KBD-GFP or Ensc-GFP) to 5 days (MBD-GFP or *ensc*), anesthetized and their ovaries were removed from the abdomen with fine forceps. Stage 10 oocytes were isolated with two 27-gauge syringe needles in Halocarbon oil (Sigma), mounted and observed by light microscopy using a DMRXA2 microscope (Leica). Velocity analyses of the particles were performed as described in two separate studies (^20, 21^) except that three kymographs were obtained for each oocyte along active flows and 5 particles (15 per oocyte) were analyzed for each kymograph using the Multi Kymograph plugin for Fiji.

### Immunofluorescence in fly oocytes

Oocytes were collected from two-day-old females. To visualize the Mt network and GFP-tagged fusion proteins, oocytes were permeabilized at 25°C in 1% Triton X100 in BRB80 buffer (80 mM K-PIPES, 1 mM MgCl_2_, 1 mM EGTA, pH 6.8) and fixed with cold methanol as described^22^. For Staufen and Gurken localization, 4-8 ovaries were fixed with PBS buffer containing 4% paraformaldehyde and 0.1% Triton X-100, washed 3 times 5 min in PBST (PBS + 0.1% Triton X-100) and blocked on PBST containing 1% BSA. Incubation with the primary antibodies was performed overnight at room temperature in BBT (PBS containing 0.1% BSA and 0.1% Tween 20). The ovaries were then briefly washed 3 times and 3 times for 30 min each in BBT. The Alexa-conjugated secondary antibody was incubated for 2 h at room temperature. The ovaries were then washed 3 times for 15 min each time in PBST, dissected, and mounted on ProLong Gold (Invitrogen).

### Protein Expression and Purification

Tubulin was isolated from porcine brain using the high-molarity PIPES procedure as previously described^23^. For bacterial expression of sfGFP-Ensconsin proteins, BL21-RIPL cells were grown at 37°C until ~O.D. 0.6 and protein expression was induced with 0.1 mM IPTG. Cells were grown at 20°C for 4 hours, harvested, and frozen. Cell pellets were resuspended in lysis buffer (50 mM Tris pH 8, 150 mM K-acetate, 2 mM Mg-acetate, 1 mM EGTA, 10% glycerol) with protease inhibitor cocktail (Roche), 1 mM DTT, 1 mM PMSF, and DNAseI. Cells were then passed through an Emulsiflex press and cleared by centrifugation at 23,000 x *g* for 20 min. Clarified lysate was passed over a column with Streptactin Superflow resin (Qiagen). After incubation, the column was washed with four column volumes of lysis buffer, then bound proteins were eluted with 3 mM desthiobiotin (Sigma) in lysis buffer. Eluted proteins were concentrated on Amicon concentrators and passed through a superose-6 (GE Healthcare) gel-filtration column in lysis buffer using a Bio-Rad NGC system. Peak fractions were collected, concentrated, and flash frozen in LN2. Protein concentration was determined by measuring the absorbance of the fluorescent protein tag and calculated using the molar extinction coefficient of the tag. The resulting preparations were analyzed by SDS-PAGE.

### Microtubule Co-sedimentation Assay

Microtubules were prepared by polymerizing 25 mg/mL of porcine tubulin in assembly buffer (BRB80 buffer supplemented with 1mM GTP, 1mM DTT) at 37°C for 15 min, then a final concentration of 20 μM taxol was added to the solution, which was incubated at 37°C for an additional 15 min. Microtubules were pelleted over a 25 % sucrose cushion at 100,000,*g* at 25°C for 10 min, then resuspended in BRB80 buffer with 1mM DTT and 10 μM taxol. Binding reactions were performed by mixing 500 nM of sfGFP-Ensconsin-KBD and/or mScarlet-K560 (that had been pre-centrifuged at 100,000g) with 2.5 μM of microtubules in assay buffer (50 mM Tris pH 8, 50 mM K-acetate, 2 mM Mg-acetate, 1 mM EGTA, 10% glycerol and supplemented with 1 mM DTT, 10 μM taxol, and 0.01 mg/mL BSA) and incubated at 25°C for 10 min. The mixtures were then pelleted at 90,000,*g* at 25°C for 10 min. Supernatant and pellet fractions were recovered, resuspended in sample buffer, run on an SDS-PAGE gel, then transferred to a PVDF membrane using an iBlot2 (Thermo Fisher Scientific) for Western blot analyses.

### Total internal reflection fluorescence microscopy (TIRF-M)

TIRF-M experiments were performed as previously described^24^ A mixture of native tubulin, biotin-tubulin, and fluorescent-tubulin purified from porcine brain (~10:1:1 ratio) was assembled in BRB80 buffer (80mM PIPES, 1mM MgCl_2_, 1mM EGTA, pH 6.8 with KOH) with 1mM GTP for 15 min at 37°C, then polymerized MTs were stabilized with 20 μM taxol. Microtubules were pelleted over a 25% sucrose cushion in BRB80 buffer to remove unpolymerized tubulin. Flow chambers containing immobilized microtubules were assembled as described^24^. Imaging was performed on a Nikon Eclipse TE200-E microscope equipped with an Andor iXon EM CCD camera, a 100X, 1.49 NA objective, four laser lines (405, 491, 568, and 647 nm) and Micro-Manager software^25^. All experiments were performed in assay buffer (30 mM Hepes pH 7.4, 150 mM K-acetate, 2 mM Mg-acetate, 1 mM EGTA, and 10% glycerol) supplemented with 0.1mg/mL biotin-BSA, 0.5% Pluronic F-168, and 0.2 mg/mL *κ*–casein (Sigma). All Kinesin-1 recruitment assays were performed in BRB80 buffer (80 mM PIPES pH 6.8, 1 mM MgCl_2_ and 1 mM EGTA) supplemented with 1 mM ATP, 150 mM K-acetate, with 0.1mg/mL biotin-BSA, 0.5% Pluronic F-168, and 0.2 mg/mL *κ*–casein. Ensconsin proteins were flowed in first, followed by K560. For live imaging, images were taken every 10 seconds for a total of 10 minutes. For fluorescence intensity analysis, ImageJ was used to draw a line across the microtubule of the K560 channel and the integrated density was measured. The line was then moved adjacent to the microtubule of interest and the local background was recorded. The background value was then subtracted from the value of interest to give a corrected intensity measurement.

### Statistical Analysis

All statistical tests were performed with Wilcoxon test or with two-tailed Student’s t-test for TIRF analyses.

## Results

### *In vivo* functional analysis of the different domains of Ensconsin

To investigate the roles of the Ensconsin functional domains (microtubule-binding and kinesin-binding), we generated several fly lines. These expressed full-length Ensc-GFP, Ensc-MBD-GFP, or Ensc-KBD-GFP. Under the control of the poly ubiquitin promoter, these lines were then used for rescue experiments in *ensc* mutant flies (Figure 1A, and Materials and Methods). We used Western blots to monitor the expression of the GFP-tagged Ensconsin variants in brain extracts. The exogenous full-length Ensc-GFP appeared as 150-kDa bands in Western blots, and it was expressed at levels similar to those of endogenous Ensconsin (Figure 1B, middle panel, lanes 6-7). The MBD-GFP (Figure 1B, top panel, lanes 2-3), and KBD-GFP proteins (lane 5), appeared as 60- and 70-kDa bands, respectively, and were expressed at similar levels as the Ensc-GFP (lane 7). In *ensc* mutant flies, Ensc-GFP expression rescued the partial lethality and restored female fertility. However, the expression of Ensc-MBD-GFP as tested in three independent transgenic lines failed to rescue the semilethality. In contrast, in two independent lines, KBD-GFP expression in *ensc*-null mutant flies rescued the partial lethality as much as wild-type (WT) Ensc-GFP, which suggests that this domain is sufficient for fly development into adulthood. However, we noticed that the Ensc-KBD flies remained sterile, suggesting that the Kinesin-binding domain alone is not enough to fully restore some aspects of oocyte development.

**Figure 1.**
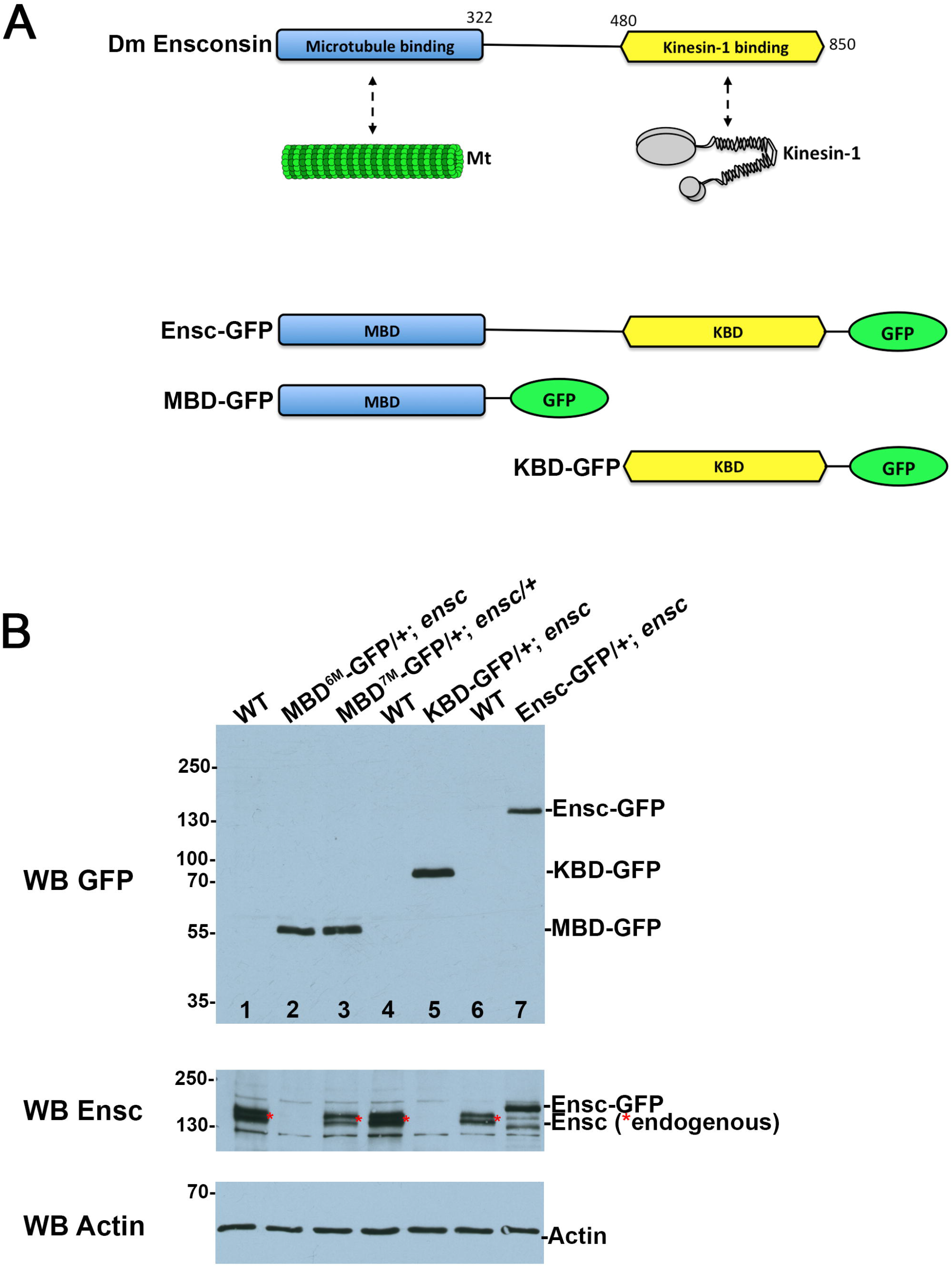
Schematic diagram and expression of GFP-tagged Ensconsin variants. A) Endogenous Ensconsin harbors an N-terminal Mt-binding domain (MBD, orange) and a C-terminal Kinesin-binding domain (KBD, blue). The 3 proteins were tagged with GFP and expressed in *ensc*-null flies using a poly ubiquitin promoter. B) Western blot analysis of different GFP-tagged Ensconsin variants from brain tissue. The membrane was probed with GFP antibodies for Western blotting (top). The same membrane was subsequently stripped (see Methods) to reveal endogenous Ensconsin (middle) and an Actin loading control (bottom). WT: control brain tissue (lanes 1, 4, 6). The expression of the transgenes is shown in ensc-null individuals (lanes 2, 5, 7-8), except in lane 3 where MBD-GFP is examined in hemizygous *ensc* tissue. The endogenous Ensconsin band is indicated with a red asterisk.

### Ensc-KBD fully rescues Kinesin-dependent centrosome separation defects in fly neuroblasts

A previous study has shown that Ensconsin stimulates microtubule growth during mitosis. As a consequence, *ensc* mitotic spindles are 20% shorter than their WT counterparts in neural stem cells (neuroblasts, Nbs)^17^. In addition, overexpression of Ensconsin (Ensc OE) in brain neuroblasts triggers the formation of longer and bent mitotic spindles (Supplementary figure 1). The fly Nb divides asymmetrically to generate a renewing Nb and a smaller cell subjected to differentiation. In this cell type, centrosome separation is initiated after the mother centrosome loses its Mt nucleation potential and is inherited in the differentiating cell, while the new centrosome retains Mt nucleation and stays in the renewing Nb^17,18,26–28^. Ensconsin and Kinesin-1 are both required during interphase to promote centrosome separation in Nbs. In *ensc* and *khc* mutants, the centrosome separation does not occur after cytokinesis, but instead just before mitosis, leading to frequent mis-segregation of mother and daughter centrosomes^17^. Interestingly, *khc* flies have normal-sized spindles, indicating that the mitotic spindle defects observed in *ensc* mutants occur independently of Kinesin-1^17^. In parallel, we measured the angles between the two centrosomes shortly before the nuclear envelope breakdown (NEBD), as readout for centrosome separation. In agreement with cooperation between Kinesin-1 and Ensconsin during interphase, we found that elevating the levels of Ensconsin protein (Ensconsin Overexpression: Ensc OE) completely rescued the centrosome separation defects in *khc* mutant neuroblasts. By contrast, *khc* mutations did not trigger spindle shortening of the long and bent spindles obtained following Ensc OE, validating that Kinesin-1 does not control spindle size in brain NBs (Supplementary Figure S1, and supplementary Video 1-4).

We therefore monitored the ability of GFP-tagged Ensconsin variants to rescue the short spindles and the centrosome separation defects observed in *ensc* mutants. To do this, we used live microscopy on Ensc-GFP, KBD-GFP, and MBD-GFP variants co-expressing RFP-tubulin. In control Nbs (Figure 2A, F-H, Supplementary Video 5), the centrosomes were fully separated before NEBD, as the angle between the two centrosomes and the nucleus center was between 120 and 180° (100%, *n* = 30). In *ensc* mutants (Figure 2B, G-H), most cells displayed incomplete centrosome separation before mitosis, as most of the angles between the two centrosomes and the nucleus center were between 60 and 120° (61.9%, *n* = 21), but this defect was restored by Ensc-GFP expression, after which 100 % (*n* = 19) of the Nbs showed pre-NEBD centrosome separations that were close to normal (Figure 2C, G-H Supplementary Video 6 and 9). Nbs expressing MBD-GFP (58.8%, *n* = 17,) showed centrosome separation defects similar to those of the *ensc* mutants (Figure 3D Supplementary Video 7). In those Nbs expressing KBD-GFP, centrosome separation was restored (87%, *n* = 23), suggesting that this domain can efficiently rescue the centrosome separation defects on its own (Figure 2E, G-H, Supplementary Video 8). Interestingly, this rescue of centrosome-separation defect did not require microtubule binding (Figure 2E, middle panels).

**Figure 2.**
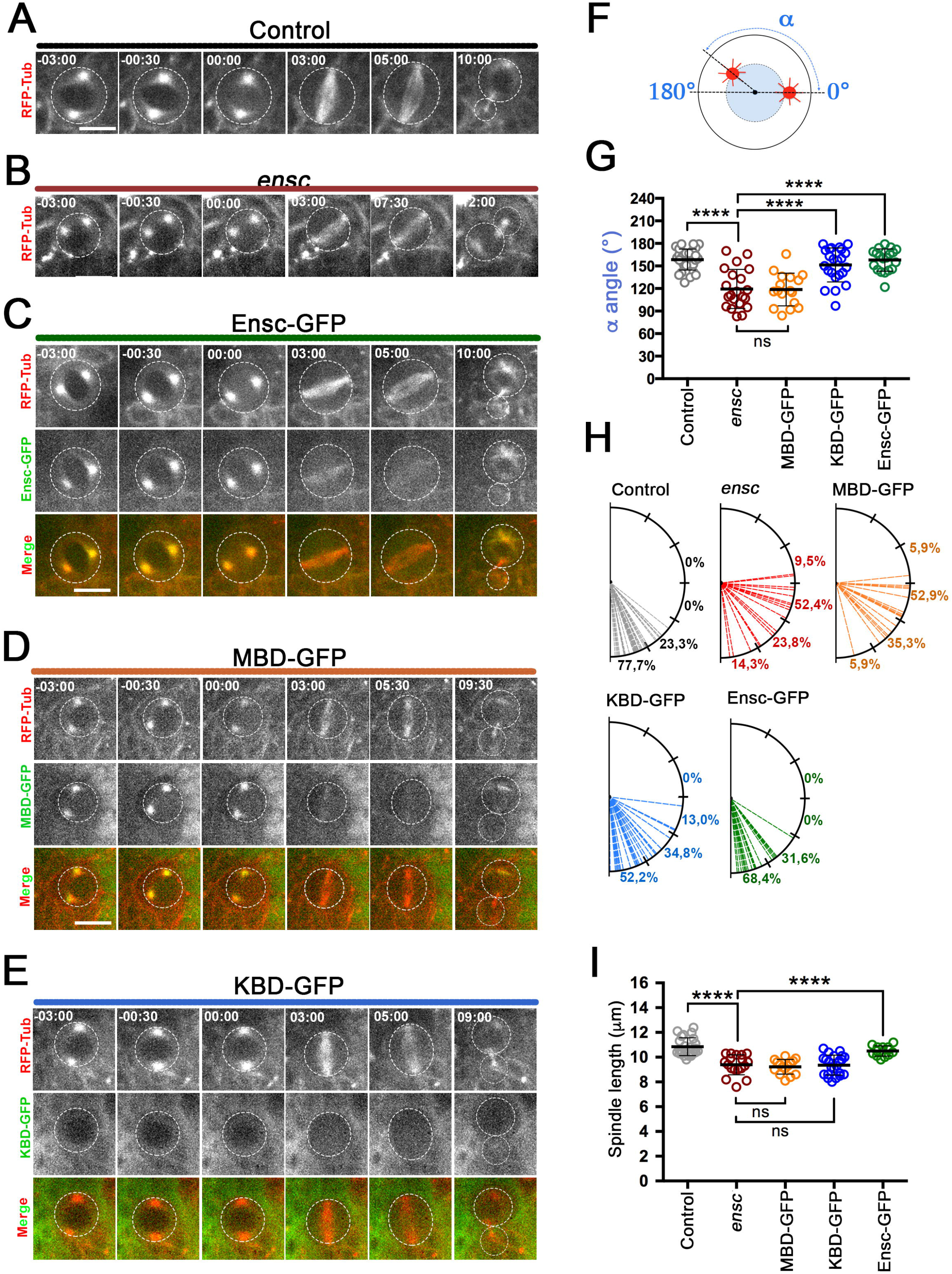
Functional analysis of microtubule- and Kinesin-binding domains during cell division. A) Selected frames of a control neuroblast and B) a mutant *ensc* Nb expressing RFP-tubulin during cell division. Three min after the nuclear envelope breakdown (NEBD), the centrosomes are already fully separated and well positioned in WT cells, but not in *ensc*. C) *ensc* mutant Nb expressing Ensc-GFP during cell division. In the merge pictures in C-E, tubulin (tub) is red and GFP-tagged proteins are green, while they are both monochrome in the upper and middle panels, respectively. D) *ensc* mutant Nb expressing MBD-GFP during cell division. Here, the centrosomes are not separated before NEBD. E) *ensc* mutant Nb expressing KBD-GFP during cell division. The centrosome separation defect is fully restored. Scale bars: 10 μm. F) α angle measurements, reflecting centrosome separation for the indicated genotypes. G) Box plot (± s.d) showing the mean centrosome separation α angle for control (158.5 ± 14.0°, *n*=30), *ensc* (119.4 ± 26.0°, *n*=21), Ensc-GFP (157.9 ± 14.9°, *n*=19), MBD-GFP (118.6 ± 21.8°, *n*=17), and KBD-GFP (151.3 ± 22.6°, *n*=23) Nbs. H) Representation of the α angle distributions of the Nbs shown in panel G. I) Box plot (± s.d) showing the mitotic spindle length for control (10.9 ± 0.7 μm, *n*=24), *ensc* (9.4 ± 0.8 μm, *n*=18), Ensc-GFP (10.5 ± 0.4 μm, *n*=13), MBD-GFP (9.2 ± 0.6 μm, *n*=15), and KBD-GFP (9.3 ± 0.8 μm, *n*=22) Nbs. ****, *P* < 0.0001 (Wilcoxon test).

Efficient mitotic spindle assembly was indicated by spindle length measurements. Ensc variant rescue of mitotic spindle length associated with Ensconsin deletion indicated that only Ensc-GFP, and not Ensc-KBD or Ensc-MBD, was able to restore normal spindle size. This demonstrates that mitotic spindle length control in neuroblasts requires full-length Ensconsin, and that MBD-GFP and KBD-GFP are unable to restore correct spindle length individually (Figure 2I, and Supplementary Video 5-6).

### The Ensconsin KBD is not sufficient for ooplasmic streaming and the correct targeting of Staufen and Gurken in the oocyte

A previous study has shown that Ensconsin is required for Staufen and Gurken transport in the oocyte to the posterior and the antero-dorsal regions, respectively^9^. We therefore monitored the localization of these two proteins in *ensc* stage-9 oocytes expressing Ensc-GFP, KBD-GFP, or MBD-GFP (Figure 3). In WT (*n*=14) and Ensc-GFP oocytes (*n*=18), Gurken was tightly localized at the cortical antero-lateral corner (Figure 3A, left, and Supplementary Figure S2). However, in *ensc* (*n*=23) and Ensc-MBD (*n*=15), most oocytes showed the Gurken protein in a punctiform pattern spread around the nuclear region (Figure 3B-C, left). KBD-GFP oocytes (*n*=23) had an intermediate phenotype: Gurken showed a WT distribution (21/23) or a partial localization at the lateral corner (2/23), with the punctiform distribution restricted to this corner (Figure 3D, left, and Supplementary Figure S2 A, B).

**Figure 3.**
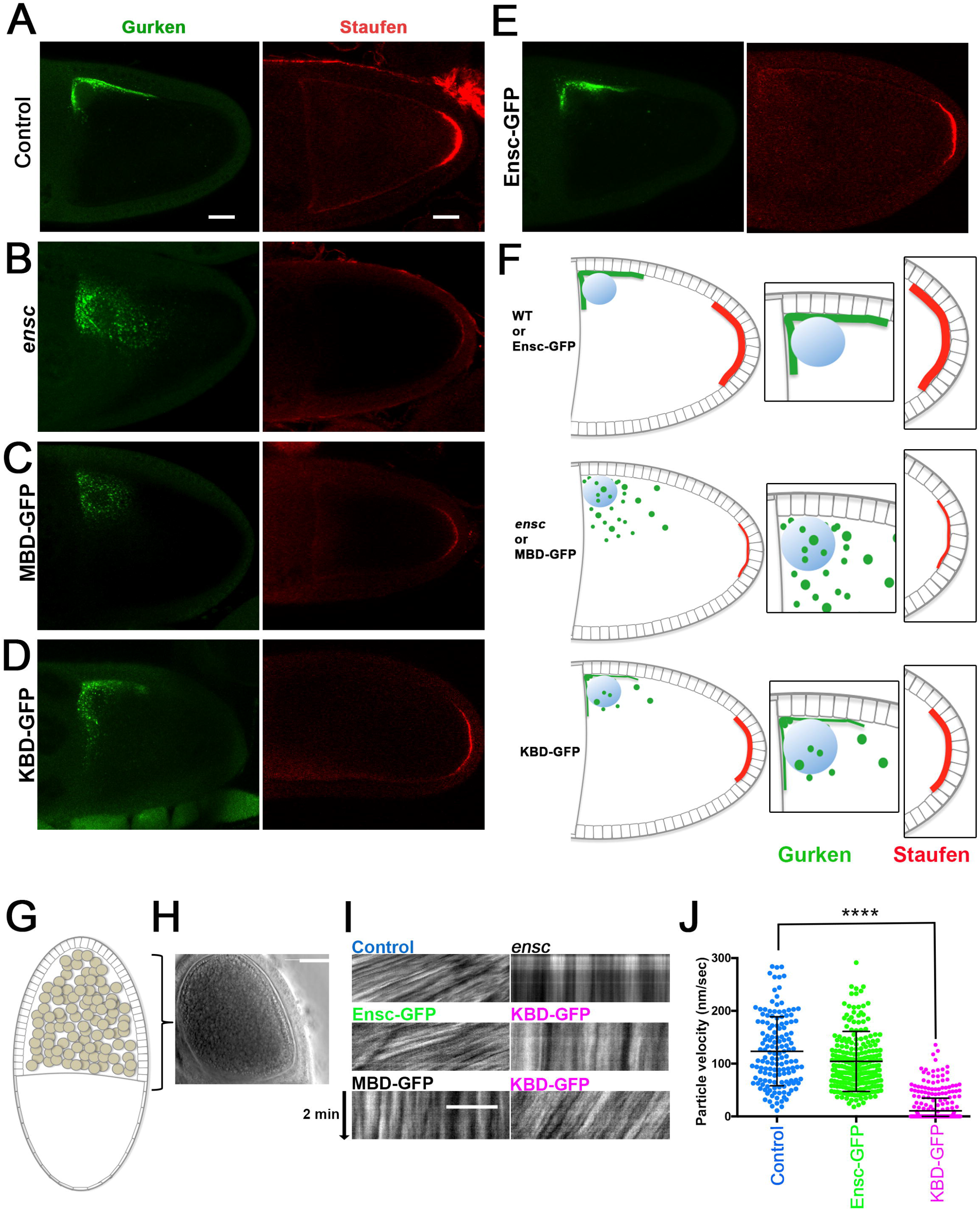
The Kinesin-binding domain of Ensconsin is not sufficient to restore the Staufen and Gurken localizations and the fast oocyte streaming in the oocyte egg chamber. A-E) Stage-9 control, *ensc*, and *ensc* mutant fly oocytes expressing MBD-GFP, KBD-GFP, or full-length Ensconsin-GFP were stained for Gurken (left, green) or Staufen (right, red). Gurken localizes at the dorsal corner side of the nucleus in both controls (*n* = 21, 100%) and in Ensc-GFP (*n* = 32). In *ensc* (*n* = 25, 100%) and MBD-GFP (*n* = 14), Gurken signal was diffused or appeared in a “spotty” diffuse pattern around the nucleus. In KBD-GFP oocytes, a fraction of Gurken correctly localized at the dorsal corner but a pool of the protein displayed a punctiform distribution at the dorsal side corner region (*n* = 41, 100%). Staufen molecules formed a robust crescent at the posterior pole of the oocyte in WT (*n* = 26, 100%) and Ensc-GFP oocytes (*n* = 29), but it was hardly distinguishable in *ensc* (*n* = 21, 100%) or MBD-GFP oocytes (*n* = 17, 100%). The KBD-GFP oocyte displayed a slightly lower Staufen crescent intensity than WT (*n* = 36, 100%). Scale bar is 20 μm. F) Schematic diagram of Gurken (green) and Staufen (red) localizations in stage-9 oocytes for WT and Ensc-GFP (top); *ensc* and MBD-GFP (middle); and KBD-GFP (bottom). Enlargements show the nucleus dorsal corner sides (middle column) with the nucleus in blue, and the embryo posterior poles (right column). G) Scheme of an oocyte. H) Phase contrast image of a control oocyte egg chamber. Scale bar: 50 μm. I) Kymographs showing oocyte streaming in different eggs chambers for 2 minutes. Scale Bar: 10 μm. The genotype is indicated at the top of each image. Control (*n*=11, Top left panel) and Ensc-GFP (*n*=16, middle left panel) expressing oocytes show similar particle velocities. Note that KBD (*n*=23) oocytes show either absent (17/23, medium right panel) or weaker ooplasmic streaming (6/23, bottom right panel). MBD-GFP oocytes (*n*=15, bottom left panel), or *ensc* oocytes (top right panel) do not show any streaming. J) Scatter dot plot showing the mean (± s.d.) particle velocities of control (*n*=146, 123±65 nm/sec), Ensc-GFP (104±56 nm/sec, *n*=243) or KBD-GFP (*n*=395, 10±23 nm/sec) ooplasms. ****: *P*<0.0001, Wilcoxon test.

At the same time, we also monitored Staufen’s localization at the posterior cell cortex. We found that Ensc-GFP (*n*=20) was able to fully rescue Staufen’s cortical posterior localization (Figure 3A, right, E, right and Supplementary Figure S2). In contrast, both in *ensc* (*n*=14) mutant oocytes and in those where MBD-GFP was expressed (*n*=13), Staufen displayed a crescent intensity 5 times weaker than that of WT (*n*=11) and Ensc-GFP oocytes (Supplementary Figure S2). Surprisingly, expression of Ensc-KBD in *ensc* mutant oocytes (*n*=13) allows Staufen to be targeted to the posterior pole in stage-9 oocytes, and the crescent intensities reached 80% of the control values (Figure 2D, right, Supplementary Figure S2). We also monitored the ability of KBD to restore ooplasmic streaming in oocyte (Figure 3, G-I) a function that is Ensconsin and Kinesin-1 dependent^9, 21, 29^. We found, as described before that *ensc* oocytes showed no ooplasmic streaming (*n*=15, Figure I). However, expression of Ensc-GFP efficiently restored ooplasmic particle velocity to near control values (Figure 3, I and J). KBD-GFP expressing oocytes showed a complete absence (17/23) or very weak (6/23) ooplasmic streaming.

Altogether, these results suggest that Ensc-KBD expression only partially restores ooplasmic streaming and the correct localization of Staufen and Gurken in *ensc* oocytes. Interestingly, this incomplete complementation seemed to be independent of Ensconsin’s microtubule binding activity: unlike MBD-GFP and Ensc-GFP, KBD-GFP did not localize to the microtubule network in the oocyte or in early dividing embryos, consistent with its lack of colocalization in S2 cells and Nbs^17^ (Figure 2 D and Supplementary Figures S3).

### The Ensconsin KBD promotes recruitment of Kinesin-1 on the oocyte Mt network

Kinesin-1 can bind to Ensconsin, and the correct targeting/loading of this motor on Mts is essential for Kinesin-1^15^. In WT stage-10 oocytes (*n*=10), Kinesin-1 heavy chain (KHC) was bound strongly to the Mt network and was easily detected by indirect immunofluorescence using commercially available polyclonal antibodies, similar to previous observations (^22^ and Figure 4A). On the other hand, KHC localization was absent on the Mt network of *ensc* oocytes (*n*=7, Figure 4, second panels from top). This localization was fully restored by the expression of Ensc-GFP (*n*=15, Figure 4, bottom panels), but not by MBD-GFP (*n*=13 Figure 4, third panels from top). In KBD-GFP-expressing oocytes (*n*=9), the Mt network showed a weak presence of Kinesin-1 (Figure 4, fourth panels from top). Altogether, these results suggest that KBD is sufficient to fully rescue centrosome-separation defects in brain Nbs and to minimally recruit of Kinesin-1 to the oocyte cytoskeleton, but can only restore partial targeting of Staufen and Gurken in the *Drosophila* oocyte.

**Figure 4.**
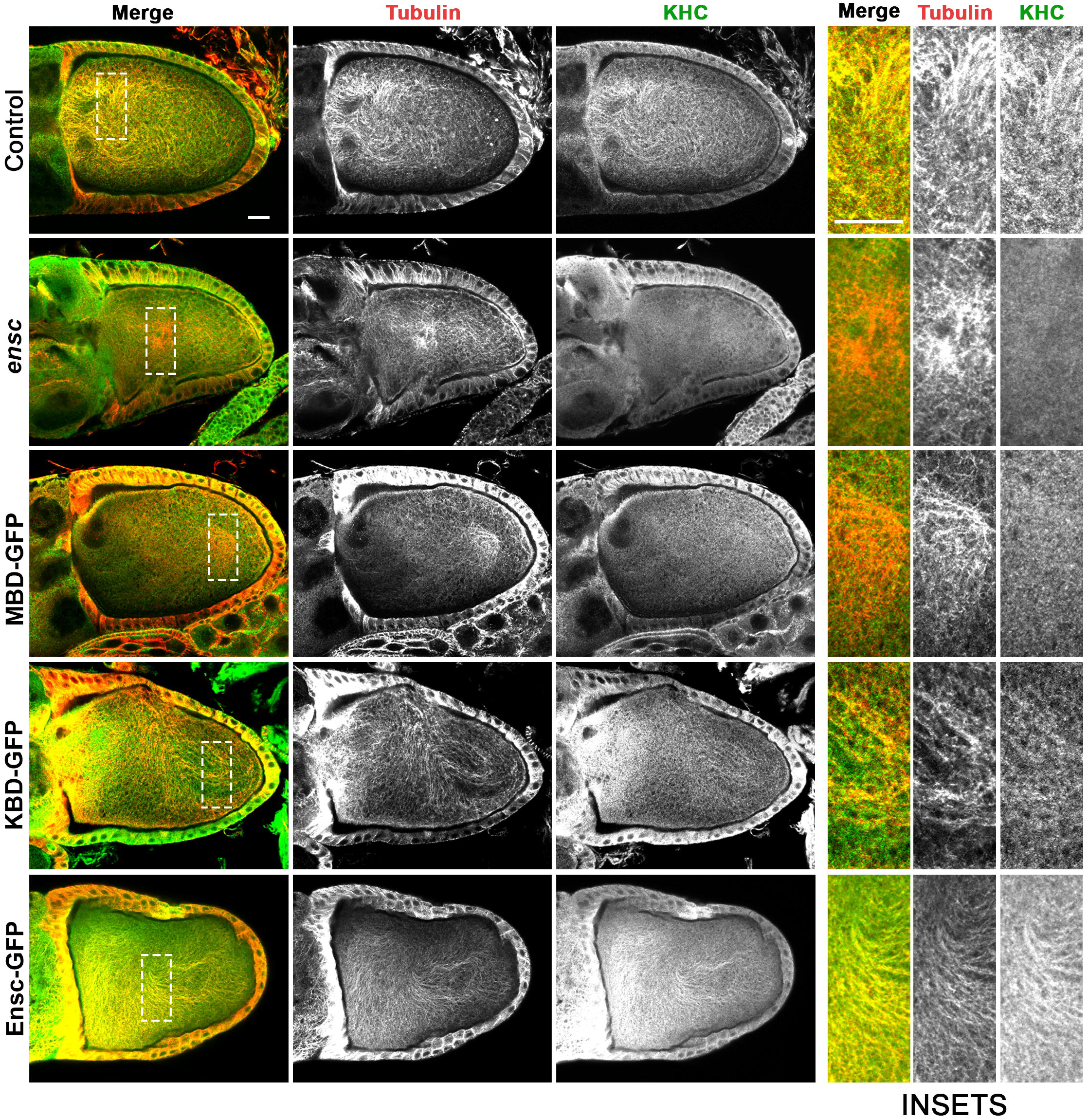
The Ensconsin Kinesin-binding domain can promote minimal recruitment of Kinesin-1 on microtubules in oocytes. Stage 10 oocytes of the indicated genotypes were permeabilized 1 hour, fixed and labeled with an anti-tubulin antibody (red in the merge column, monochrome elsewhere) or an anti-Kinesin-1 one (green in the merge column, monochrome elsewhere). Scale bar = 20 *μ* m. WT (n=10) and Ensc-GFP (n=15) shows a strong KHC labeling on the Mts, by contrast to *ensc* (n=7) or MBD-GFP (n=13) oocytes where KHC is not detectable. The right part of the image shows an enlargement view of the Mt network. KBD-GFP oocytes display a moderate recruitment of KHC (n=9).

### The Ensconsin KBD modestly recruits Kinesin-1 to Mts *in vitro*

To assess whether the KBD recruits Kinesin-1 on Mts directly, we used an *in vitro* approach with purified components (Figure 5). Total internal reflection fluorescence (TIRF) microscopy allowed us to monitor and quantify the recruitment of Kinesin motor protein molecules on stabilized Mts in the absence or presence of Ensconsin variants. Although full-length Ensconsin recruited the truncated Kinesin motor, K560, 25-fold to the microtubule, the KBD did not significantly recruit K560 above K560 alone in our conditions (Figure 5A-C). However, this assay uses very low concentrations of proteins and therefore a modest recruitment may not be apparent. We therefore performed a co-sedimentation assay with purified proteins and taxol-stabilized microtubules to analyze the effect of KBD on K560 recruitment in an excess of Mts. Interestingly, in these conditions; we found a modest enrichment of K560 in the Mt pellet in the presence of KBD compared to the absence (Figure 5D). In addition, we found that KBD was apparent in the Mt pellet only in the presence of K560, but not in its absence, suggesting these two proteins may form a complex that exhibits a higher affinity for Mts. Altogether, our data indicate that KBD is able to recruit enough Kinesin-1 to Mts both *in vitro* and *in vivo* to rescue the defects observed in the absence of Ensconsin, depending on cellular context.

**Figure 5.**
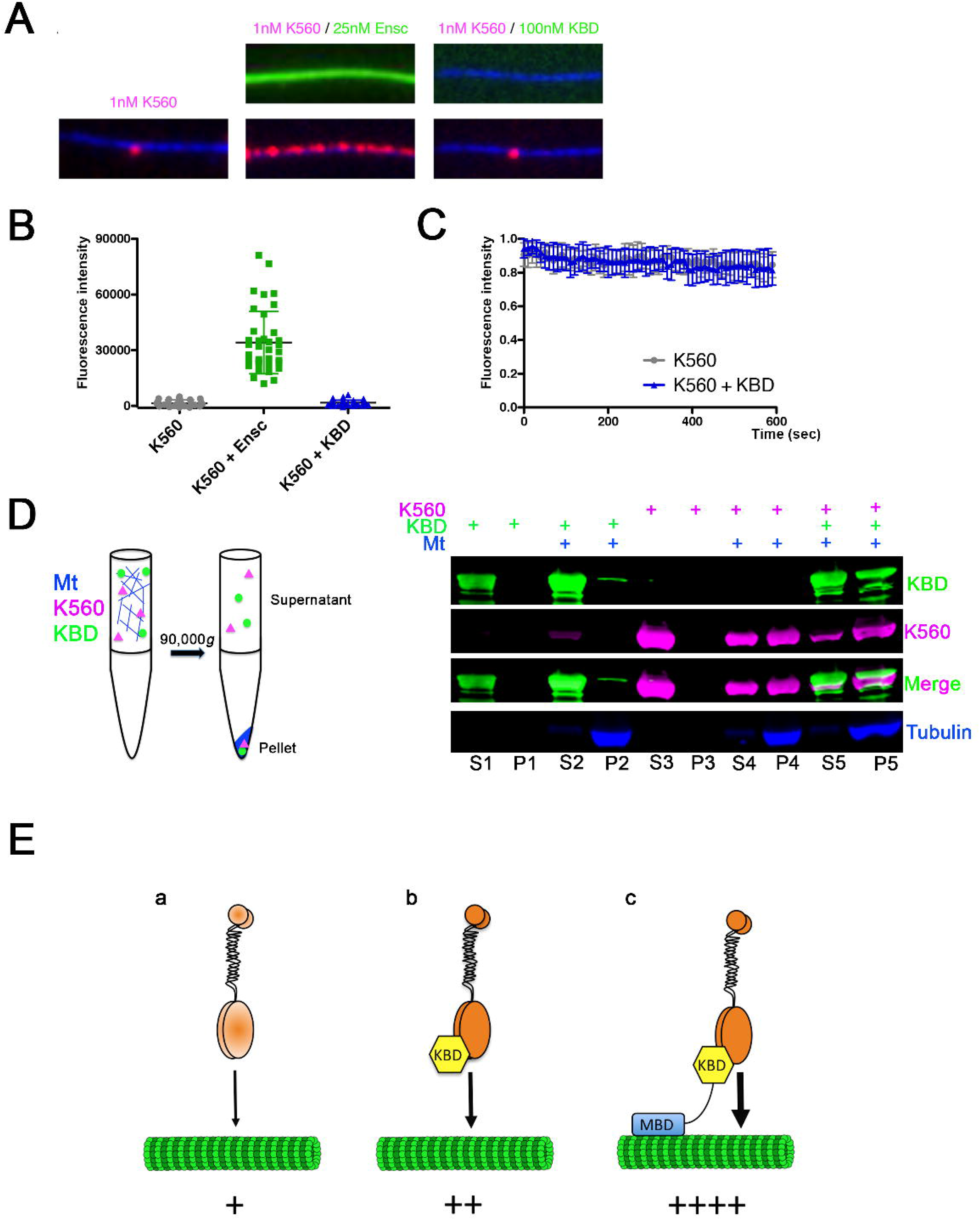
The Ensconsin Kinesin-binding domain can promote modest recruitment of K560 to Mts *in vitro*. A) TIRF-M images and corresponding kymographs of 1 nM K560-mScarlet (magenta) + 1 mM ATP in the absence and presence of 25 nM sfGFP-Ensc or 100 nM sfGFP-KBD (green). Images are 7.0 μm in length. B) Quantification of fluorescence intensity of K560-mScarlet in the absence and presence of 25 nM sfGFP-Ensc or 100 nM sfGFP-KBD (means ± s.d. are 1322.4 ± 1889.7, 34160.3 ± 16816.5, and 1668.8 ± 1410.3 A.U., and *n* = 33, 38, and 37 microtubules for K560 alone, K560 + Ensc, and K560 + KBD, respectively from two independent trials; *P* < 0.0001 for K560 vs. K560 + Ensc, and *P* = 0.385 for K560 vs. K560 + KBD, Student test). C) Quantification of K560 fluorescence intensity in the absence and presence of KBD over 10 min at 10 s intervals (*n* = 12 and 14 microtubules for K560 and K560 + KBD, respectively, quantified from two independent trials). D) Left: Scheme of the Mt binding assay. KBD and K560 were mixed in the presence of polymerized Mts and subjected to centrifugation. Pellets (P) and Supernatants (S) were subsequently analyzed by Western blot. Right: Western blot shows the binding behavior of 500 nM Ens-KBD (green) and/or 500 nM K560 (pink) in the presence of 2.5 μM taxol-stabilized microtubules (blue). See the increase of K560 in the Mt pellet in the presence of KBD (purple, compare P4 with P5 and S4 with S5). Note that KBD is only detected in the Mt pellet in the presence of K560 indicating direct interaction (green, compare P5 with P2). E) Model for Kinesin-1 recruitment to Mts. Panel a: Kinesin-1 (light orange) shows intrinsic Mt binding. Panel b: Kinesin-1 intrinsic binding to Mts can be stimulated (dark orange) after interaction with KBD (yellow). Panel c: *in vivo*, the stimulation of Kinesin-1 binding to Mts occurs on the Mt directly on the Mt and is very efficient.

## Discussion

### Full-length Ensconsin/MAP7 is required for maintaining mitotic spindle size

Ensconsin/MAP7 was first discovered two decades ago and was originally purified from microtubule pellets isolated from epithelial cells^5^. Preliminary overexpression experiments did not reveal modified microtubule dynamics in human interphase cells, and *ensc* mutant oocytes have normal Mt cytoskeletons^6^. However, changes in Ensc/MAP7 levels affect spindle morphogenesis in fly neural stem cells and in human cultured cells, and a recent study showed that human Ensconsin removal from Mts is crucial at NEBD to prevent spindle defects^17, 30, 31^. Fly *ensc* mutant Nbs display short spindles and have reduced Mt polymerization speeds. In addition, Ensconsin strongly stimulates Mt polymerization *in vitro*, and its overexpression increases spindle length (Unpublished results and Supplementary Figure S1). Rescue experiments indicate that neither MBD-GFP nor KBD-GFP fusion proteins are able to restore spindle length, even when expressed at similar levels to full-length Ensconsin-GFP, which efficiently rescues that phenotype. This reveals that although it has a functional Mt-binding sequence, the MBD is not able to stimulate Mt polymerisation on its own *in vivo*. These results are in agreement with the observation that in *Drosophila* S2 cells, unlike with the overexpression of the full-length protein, neither the overexpression of the KBD or of the MBD alone can lead to increases in mitotic spindle length (unpublished and^17^). Further studies at the molecular level will be needed to investigate how the MBD and KBD domains cooperate to stimulate Mt polymerisation *in vitro*.

### Mechansim of Ensconsin/MAP7 stimulation of Kinesin-1 targeting to Mt

Several studies have shown a strong connection between Kinesin-1 and Ensconsin. First of all, inhibition of Ensconsin and Kinesin-1 has yielded similar phenotypes in four different studies^9, 15–17^ Secondly, we know that Ensconsin and the Kinesin motor domain can interact *in cellulo*. However, this interaction has only been seen with simultaneous overexpression of both the KBD and the Kinesin motor domains, which suggests that the interaction between the two is weak and/or transient^15, 32^ This transience is likely critical because efficient transport would be hurt by a strong attachment to Mt-bound Ensconsin, which has high salt-resistant interactions with the Mts^5^. Congruently, Kinesin and Ensconsin single motors have never been seen to move together with Ensconsin molecules *in vitro*^32^

Based on the current literature, there are two models for Kinesin-1 activation/recruitment by Ensconsin/MAP7. In the first one, Ensconsin bound (via the MBD) to the Mt lattice serves as a direct recruitment platform for Kinesin-1 (Figure 5). This targeting model is supported by Kinesin-1 motor recruitment on Mts, which is lower in *ensc* oocyte extracts and undetectable in intact *ensc* oocytes (^9^, this study, and Figure 4). Furthermore, in Ensconsin and Kinesin-1 knockdown muscle cells, a fusion protein between the Ensc-MBD and Kinesin-1 motor domain (leading to strong recruitment of Kinesin motor to the Mt network) was able to rescue nuclei spacing, thus strongly supporting that the function of Ensconsin is to deliver Kinesin-1 to Mts in large cells such as muscles and oocytes^15^. Finally, this direct recruitment platform model is supported *in vitro*, as the Kinesin-1 motor is strongly recruited on Ensconsin-labeled Mts (^32^ and Figure 5).

In a second model based on S2 cells studies, the Ensc-KBD is necessary and sufficient to promote full Kinesin-1 activation^16^. This study demonstrates that this activation mode is not sufficient *in vivo*. In the fly Nb, both Ensconsin and Kinesin-1 participate in the centrosome separation process^17^. As we show here, this defective centrosome separation phenotype is completely restored by the KBD. By contrast, this rescue is not validated in *ensc* mutant oocytes. In the absence of Ensconsin, ooplasmic streaming is abrogated and asymmetric transport towards the oocyte’s posterior cortex (Staufen) and toward the dorsal corner (Gurken) is impaired. We found that KBD expression promotes only partial Gurken and Staufen localization but not efficient oocyte streaming, but is less efficiently than when carried out by the full-length MBD-containing Ensconsin. Remarkably, this partial rescue occurs with very weak recruitment of Kinesin-1 on the Mts (at least when measured using conventional techniques), thus reinforcing the idea that a low amount of Mt-bound Kinesin-1 molecules is enough to drive minimal transport in the oocyte.

How does the binding of KBD to Kinesin-1 support its activation? Interestingly, we show here for the first time that KBD forms a protein complex with the Kinesin-1 motor to stimulate its binding to Mts (Model figure 5E, panel b). Altogether, we propose that Kinesin-1 binding to Mts is subjected to a dual regulation by Ensconsin. First, an Ensconsin-decorated Mt provides a platform to recruit Kinesin-1. Second, direct interaction with the KBD stimulates affinity of the recruited motor to the Mt (Figure 5E, panel c). We can speculate that further studies will reveal that this dual regulation of Kinesin-1 binding to Mt by Ensconsin/MAP7 is also regulated by post-translational modifications (e.g. phosphorylation) to orchestrate activation of Kinesin-1 in space and time and in different contexts.

Our data suggest that, at least in *Drosophila*, the model based on Kinesin-targeting on Mts by full length Ensconsin is not an absolute requirement for development, since KBD flies are viable, at least in the lab conditions where flies are maintained under optimal temperature and media growth conditions. However, unlike flies rescued by full-length Ensc, KBD females remain sterile possibly because *ensc* meiotic spindle are abnormal and do not support correct chromosome segregation (Personal communication, Prof. Ohkura, University of Edinburgh). In addition, the KBD could not ensure ooplasmic streaming or the optimal localization of the Kinesin-1-dependent cargoes analyzed in this study, including Gurken and Staufen in oocytes. The length of the egg chamber of a stage-9 oocyte is 120 μm, so oocytes are remarkably larger than Nbs and S2 cells, which have diameters of 10-12 μm. In addition, the oocyte egg chamber is a crowded environment that displays a very complex Mt network as compared to most fly cells^20, 22^ In this context, Kinesin-1-mediated asymmetric and bidirectional transport is a difficult task that probably requires the strong attachment of Kinesin-1 to the complex Mt network in the fly oocyte. In agreement with this hypothesis, the fly oocyte is the only cell type in which conventional immunofluorescence methods detect strong KHC labeling on Mts (this study,^22^). We therefore propose that this strong targeting may be unnecessary in most fly cell types, but that in large cells having complex Mt networks, the optimal targeting/loading of Kinesin-1 onto microtubules may be essential for optimal Kinesin-1 transport over long distances.

## Acknowledgments

This work was funded by the Ligue Nationale Contre le Cancer, l’Association de la Recherche contre le Cancer. We would like to thank R. Basto for the kind gift of RFP-tubulin expressing flies. We thank Kahina Sadaouli for the preliminary functional analyses of Ensc domains during oocyte development. We thank the Microscopy Rennes Imaging Center for the microscopy facilities. M. M. is a doctoral fellow of the Région Bretagne and the Ligue Nationale contre le Cancer. E. G. was a doctoral fellow of the Région Bretagne. R. C was a doctoral fellow of the French Ministry of Research and the Association de la Recherche contre le Cancer. AG was supported by the CNRS and by the Association pour la Recherche sur le Cancer (grant PJA 20161204931). KMOM is supported by the NIH (R00HD080981), the March of Dimes Foundation, and the Simons Foundation. We thank Christelle Benaud and Romain Gibeaux for critical reading and helpful comments on the manuscript. The authors have no competing financial interests to declare.

---

**Supplementary Figure S1. Overexpression of Ensconsin increases mitotic spindle length in a Kinesin-1-independent manner but efficiently rescues *khc*-dependent centrosome separation defects and in brain Nbs**. A) WT, Ensconsin-Venus overexpressing (Ensc OE), *khc^27^/khc^63^* and *khc^27^/khc^63^*, Ensc OE Nbs were imaged during cell division. Scale bar is 10 μm. B) Box plot showing the mean (± s.d.) mitotic spindle length analysis for control (11.3 ± 0.8 μm, *n*=14), *khc^27^/khc^63^* (10.9 ± 0.5 μm, *n*=17), Ensc OE (13.6 ± 0.9 μm, *n*=24), *khc^27^/khc^63^*, Ensc OE (13.5 ± 2.0 μm, *n*=24) Nbs. ***, *P* < 0.001 (Wilcoxon test). C) Analysis of c□□□□□□□□□ separation angle (α at NEBD. D) Box plot showing the mean (± s.d.) centrosome separation angle for control (151.4 ± 19.2°, *n*=17), *khc^27^/khc^63^* (114.1 ± 20.6°, *n*=21), Ensc OE (134.0 ± 40.3°, *n*=26), *khc^27^/khc^63^*, Ensc OE (159.4 ±16.0°, *n*=29) Nbs. ****, *P* < 0.0001 (Wilcoxon test).

**Supplementary Figure S2. Gurken and Staufen localization in oocytes.** Stage 9 oocytes from WT, *ensc*, KBD-GFP, MBD-GFP or Ensc-GFP were fixed and stained with anti-Gurken antibodies. A) Antero-posterior localization of Gurken in a WT oocyte (top); partial Gurken localization in a KBD-GFP-expressing oocyte, with some of the protein dispersed (middle); and an *ensc* mutant oocyte in which most of the Gurken protein is dispersed around the nucleus (bottom). The scale bar is 20 μm. B) Histogram showing the repartition of the Gurken protein in stage-9 oocytes: WT (*n*=14); *ensc* (*n*=14); MBD-GFP (n = 15); KBD-GFP (*n*=18); and Ensc-GFP (*n*=23). C) Staufen crescent intensity observed in control, *ensc*, MBD-GFP, KBD-GFP, and Ensc-GFP stage-9 oocytes. D) Box plot showing the normalized mean crescent intensity (± s.d.) of stage-9 oocytes for control (1.00 ± 0.33, *n*=11); *ensc* (0.33 ± 0.17, *n*=14); MBD-GFP (0.37 ± 0.14, *n*=13); KBD-GFP (0.89 ± 0.33, *n*=13); and Ensc-GFP (1.01 ± 0.20, *n*=20) transgenic lines. ****, *p* < 0.0001; ***, *p* = 0.0007 (Wilcoxon test).

**Supplementary Figure S3. Localization of Ensc-GFP, KBD-GFP, and MBD-GFP variants and of endogenous Ensconsin in oocytes and of KBD-GFP, and MBD-GFP in live embryos.** Oocytes for the indicated genotypes were permeabilized in an Mt-stabilizing buffer for 1 h to remove cytoplasmic protein pools. They were then fixed and immuno-labeled with an antitubulin antibody (red on the left, monochrome elsewhere) plus either (A) an anti-GFP antibody or (B) an affinity-purified anti-Ensconsin antibody (both antibodies are green in the left panels, monochrome elsewhere). Scale bars are 20 μm and 50 μm respectively for panels A and B. Note that KBD-GFP is highly expressed, but that the protein is removed after permeabilization indicating the protein is cytoplasmic. Endogenous Ensconsin, Ensc-GFP and MBD-GFP are found on the oocyte Mt cytoskeleton. C) Selected images of an embryo expressing MBD-GFP as it divides during interphase (at 4:30, left) and during metaphase (8:00, right). C) Selected images of a dividing embryo expressing KBD-GFP, shown during prophase (1:00, left) and metaphase (4:30, right). The GFP-tagged proteins are green and RFP-tubulin is red in the merge pictures, and they are both monochrome in the other panels. Scale bars: 20 μm. Time is indicated as min:sec. MBD-GFP co-localizes with Mts during mitosis and interphase, but KBD-GFP does not.

